# Nanoparticle coatings for controlled release of quercetin from an angioplasty balloon

**DOI:** 10.1101/2022.04.28.489902

**Authors:** Ioana Craciun, Carlos E. Astete, Dorin Boldor, Merilyn H. Jennings, Jake D. Gorman, Cristina M. Sabliov, Tammy R. Dugas

## Abstract

Peripheral artery disease (PAD) is a systemic vascular disease of the legs that results in a blockage of blood flow from the heart to the lower extremities. Now one of the most common causes of mortality in the U.S., the first line of therapy for PAD is to mechanically open the blockages using balloon angioplasty. Coating the balloons with antiproliferative agents can potentially reduce vessel re-narrowing, or restenosis after surgical intervention, but current drug-coated balloons releasing chemotherapy agents like paclitaxel have in some cases shown increased mortality long-term. Our aim was to design a novel drug-coated balloon using a polymeric nanodelivery system for a sustained release of polyphenols that reduce restenosis but with reduced toxicity compared to chemotherapy agents. Poly (lactic-co-glycolic acid) (PLGA) nanoparticles with entrapped quercetin, a dimethoxy quercetin (rhamnazin), as well as quercetin conjugated to PLGA, were developed. Balloon catheters were coated with polymeric nanoparticles using an ultrasonic method, and nanoparticle characteristics, drug loading, coating uniformity and drug release were determined. The adhesion of nanoparticles to vascular smooth muscle cells and the antiproliferative effect of nano-delivered polyphenols were also assessed. Of the nanoparticle systems tested, those with covalently attached quercetin provided the most sustained release over a 6-day period. Although these particles adhered to cells to a smaller extent compared to other nanoparticle formulations, their attachment was resistant to washing. These particles also exhibited the greatest anti-proliferative effect. In addition, their attachment was not altered when the cells were grown in calcifying conditions, and in PAD tissue calcification is typically a condition that impedes drug delivery. Moreover, the ultrasonic coating method generated a uniform balloon coating. The polymeric nanoparticle system with covalently attached quercetin developed herein is thus proposed as a promising platform to reduce restenosis post-angioplasty.

## 1. Introduction

Peripheral artery disease (PAD) is a systemic atherosclerotic disease that affects approximately 202 million people worldwide. With over 8 million diagnoses, PAD is one of the most common causes of mortality in the United States (1-4). Moreover, atherosclerotic diseases like PAD are becoming a world-wide problem (5). PAD is characterized by debilitating atherosclerotic occlusion of arteries in the lower extremities, resulting in an obstruction of blood flow (1, 6). Though a disease of the extremities, left untreated, PAD can culminate in catastrophic consequences like stroke, myocardial infarction, and death (2, 7). The most common symptom among patients with PAD is intermittent claudication, but it is often asymptomatic, under-diagnosed and under-treated, resulting in a reduced functional capacity and quality of life. In its most severe form, the resulting limb ischemia can necessitate limb amputation (2-4). To treat lower extremity PAD, clinicians often revascularize the affected artery or arteries using an endovascular procedure known as angioplasty, achieved using balloon dilation and sometimes, placement of a stent (8-11). Angioplasty is a technique of mechanically widening a blood vessel that has been narrowed or obstructed due to atherosclerosis (12). In PAD, balloon angioplasty is favored over stenting due to the small diameter of the affected arteries and the preponderance of stent fractures occurring in clinical cases (13, 14). Balloon angioplasty allows slow vessel stretching to enlarge the lumen (12). Unfortunately, it also induces stretch and strain to the vessel wall, and the injury it imparts induces a series of cellular events culminating in the formation of a new lesion (15). Restenosis or vessel re-narrowing after implantation remains a complication of vascular interventions (16). Early restenosis and neointimal hyperplasia within the stented vessels have been attributed to deep vascular injury, with fracture of the internal elastic lamina (17). Intimal hyperplasia includes inflammatory phenomena, migration, and proliferation of smooth muscle cells and also, extracellular matrix deposition (17). These events culminate in a thickened vessel wall that obstructs blood flow (15, 18).

Current protocols for the prevention and therapy of restenosis after angioplasty/stenting are based on sustained, antiproliferative drug release into the vessel wall (19). Drug-coated balloons (DCB) have recently emerged as a treatment for peripheral artery (19-22) and coronary in-stent restenosis (23). The concept of DCB therapy relies on healing of the vessel wall after a rapid release of drug locally but retention of the drug within the vessel wall long enough to impact deleterious cellular events occurring early after the procedure. DCBs require three fundamental elements: a semi-compliant angioplasty balloon, an antiproliferative drug and a drug carrier (23). DCB releasing the chemotherapy agent paclitaxel have been approved by the FDA. Paclitaxel is highly lipophilic and binds quickly and tightly to tissue, which results in rapid cellular uptake and long-term retention at the site of delivery. This treatment comes with major disadvantages such as: systemic toxicity (15, 24), the release of paclitaxel before arrival at the lesion site due to direct application of drug to the balloon surface (15, 24) and delayed re-endothelialization, as demonstrated by animal studies utilizing paclitaxel-eluting stents (25, 26). In addition, recent alerts issued by the FDA identified a late mortality signal in study subjects treated with paclitaxel-coated balloons. The relative risk for increased mortality at 5 years was 1.57 (95% confidence interval 1.16 – 2.13), which corresponds to a 57% relative increase in mortality in patients treated with paclitaxel-coated devices (27). Therefore, studies focused on controlled delivery of other anti-proliferative agents have evolved. Our own prior research focused on two synergistic polyphenols - resveratrol and quercetin - and these studies demonstrated that the two have low toxicity and reduce vascular smooth muscle proliferation but promote re-endothelialization, both *in vitro* and *in vivo* (15, 28). We were also successful in developing a drug-eluting coating that successfully achieves slow release of resveratrol (i.e., over several days), but by comparison, release of quercetin was more rapid and less protracted (15).

Within this framework, the aim of this study was to develop polymeric nanoparticles (pNP) for quercetin delivery that were capable of a high entrapment, slow release of drug and antiproliferative activity. Poly (lactic-co-glycolic acid) (PLGA) nanoparticles with entrapped quercetin (pNP(eQ)), a dimethoxy quercetin (i.e., rhamnazin, designated pNP(eR)), as well as quercetin conjugated to PLGA (pNP(cQ)), were developed. Using an ultrasonic coating method, miniaturized balloon catheters were coated with pNP, and nanoparticle characteristics, drug loading, drug release, and efficacy in reducing vascular smooth muscle cell proliferation were assessed. With respect to the coated balloons, we also determined particle deposition on the balloon surface, assessed as total pNP and drug loading, as well as coating uniformity. We aimed to achieve uniformly coated balloons, with the particles firmly adhered. Our overarching project goal is to achieve minimal loss of drug from the balloon surface during transit to the lesioned area, but upon inflation within the lesioned artery, the particles transfer and attach firmly to the vessel wall, where the coating begins releasing polyphenols. As such, we aim to achieve a controlled and localized administration of the active substance in the affected area.

A secondary aim of our design was to enable pNP adhesion to calcified lesions. Vascular calcification is a common occurrence in PAD and compared to coronary artery disease, can be extensive (29). The accumulation of calcium and phosphate in the intimal and medial layers of the vessel are typical of patients with PAD, particularly those with chronic kidney disease and diabetes mellitus (30). Calcification is a key contributor to poorer outcomes after angioplasty, as it leads to altered compliance, flow-limiting dissections and acute vessel recoil (31). Moreover, late lumen loss after paclitaxel-coated balloon therapy was shown correlated with circumferential calcification (32), and hypotheses are that such outcomes are due to an inability of the calcified lesion to absorb paclitaxel. Thus, in some experiments, we tested whether our pNP coating was capable of strong adhesion to cells in which calcium accumulation was induced experimentally.

## 2. Materials and methods

### 2.1. Materials

The following materials were obtained from Sigma-Aldrich (St. Louis, MO): Resomer RG504H poly (D, L-lactide-co-glycolide), PLGA 50:50 (molecular weight 38,000- 54,000), acetone and poly (vinyl alcohol) (PVA 31,000-50,000; 87-89% hydrolysed), quercetin and rhamnazin. PLGA covalently modified with quercetin was synthetized in the laboratory. Analytical grade chemicals and reagents were used for this study.

### 2.2. Conjugation of PLGA with quercetin

The coupling of quercetin to PLGA was based on an acylation reaction. The first step was PLGA activation. Briefly, 2 g of PLGA was dissolved in 50 mL DCM at room temperature in a 3-neck round bottom flask. A bubbler bottle with 1 M sodium hydroxide NaOH was required to neutralize HCl produced during the reaction under nitrogen. After complete dissolution of PLGA at room temperature, 10 eq. of oxalyl chloride was added dropwise with a glass syringe, along with 3 mL of DMF. The reaction was performed at room temperature with mild stirring for 5 hours. Next, the solution was concentrated with a Buchi R-300 Rotavapor (Buchi Corporation, New Castle, DE). The activated PLGA polymer was precipitated by addition of 200-300 mL of ethyl ether. The white precipitate was washed at least three times with ethyl ether to remove impurities. The solids were dried overnight under high vacuum. The second reaction was performed by dissolution of 1 g of dry PLGA-Cl in 25 mL of DMSO, which was added dropwise to 35 mg of quercetin dissolved in 20 mL of DMSO. The reaction was performed overnight at room temperature under nitrogen. The PLGA-quercetin polymer was precipitated by addition of 150 mL of ethyl ether; the precipitation was repeated three times. The precipitated polymer was suspended in 80 mL of DCM and the organic phase was washed with 200 mL of water to remove unreacted quercetin. The process was repeated to obtain a clear supernatant. Finally, the DCM was evaporated with a Buchi R-300 Rotavapor, and the polymer was dried under high vacuum for 3 days at 30°C. The PLGA-quercetin copolymer was stored at 2-4°C for further characterization and use in nanoparticle synthesis.

### 2.3. pNP synthesis

#### 2.3.1. Synthesis of PLGA-Eudragit RL-100 nanoparticles

The polymeric nanoparticles were synthesized employing a single emulsion evaporation technique. Briefly, an organic phase was created by mixing Eudragit RL 100 (60 mg) and PLGA (200 mg) in ethyl acetate to acetone (8:2) solution (6 mL), with mild stirring at room temperature for 30 minutes. Next, quercetin or rhamnazin was added to the organic phase. Rhamnazin was used to test whether alkylation of quercetin resulted in a protracted release profile. After 15 min and with continued stirring at room temperature, the organic phase was poured dropwise into 60 mL of aqueous phase containing 4 mg/mL Tween 80. To reduce droplet size, the emulsion was microfluidized with an M-110P Microfluidizer (Microfluidics Corp, Westwood, MA) at 4°C, 30,000 PSI, with four passes. Ethyl acetate in the suspension was evaporated using a Buchi R-300 Rotavapor (Buchi Corp., New Castle, DE) under vacuum at 32°C for 2 h. Finally, the nanoparticle suspension was mixed with trehalose at a 1:2 mass ratio, and the suspension was freeze-dried with a FreeZone 2.5 (Labconco Corp., Kansas City, MO) at 32°C for 2 days. A 2 mL solution of polyvinyl alcohol (PVA; 30 mg) was added before freeze-drying to minimize aggregation after polymeric nanoparticle resuspension. The powdered samples were kept at -80°C until further characterization and use. In some studies, PLGA was covalently modified with Q prior to pNP synthesis (see section 2.2), but all other steps were identical. The mean size, PDI and zeta potential of the polymeric nanoparticles were measured by Dynamic Light Scattering (DLS) with a Malvern Zetasizer nano ZS (Malvern Panalytical inc, Westborough, MA). Because pilot studies demonstrated an impact of trehalose on cell growth, for studies examining the effect of nano-delivered quercetin on vascular smooth muscle cell proliferation, the pNP were prepared fresh on the day of the experiment, without freeze-drying and without trehalose. However, all other components were maintained at a similar ratio to ensure that the pNP formulations for the two studies were similar.

### 2.4 Nanoparticle characterization

#### 2.4.1 Morphology

Transmission electron microscopy (TEM) was accomplished using a JEOL JM-1400 (JEOL USA Inc., Peabody, MA) and an accelerating voltage of 120 kV. As such, TEM was used to analyse the structure of empty PLGA polymeric nanoparticles (pNP(E)), PLGA NP with entrapped rhamnazin (pNP(eR)), PLGA pNP with entrapped quercetin (pNP(eQ)) and PLGA NP with conjugated quercetin (pNP(cQ)). One drop of the pNP resuspension in nanopure water was placed onto a carbon film 400 mesh copper grid, and the excess amount of solution was removed with sterile filter paper. A solution of 2% uranyl acetate was used for staining. After 5 min, a separate sterile filter paper was utilized to remove excess uranyl acetate.

#### 2.4.2 Size Distribution and Zeta Potential Characterization

Dynamic light scattering (DLS) (Malvern Panalytical, Westborough, MA) was employed to characterize the nanoparticles for size, polydispersity and zeta potential. After resuspension in low resistivity water, a disposable capillary cell of 1 mL volume was used to measure size, polydispersity index (PDI), and zeta potential (Smoluchowski model) for NP.

### 2.5 Drug release and biologic efficacy

#### 2.5. Drug release protocol

The release profiles were performed by placing 10mg/mL PLGA-Eudragit RL100 NP (pNP(eQ), pNP(eR), pNP(cQ)) in dialysis membrane (molecular weight cut-off of 12,000/14,000 g/mol, regenerated cellulose, Fisher Scientific). Sterile PBS was used for sample resuspension. The samples were dialyzed against 800 mL of PBS at 37°C under continuous stirring, and PBS was replaced every 8 h in the first 12h and then every 24h. At pre-determined time points, 0.2 mL samples were taken from inside the dialysis bag (nanoparticle solution) and to prevent quercetin oxidation, was mixed with 20 μL of 50 mM ascorbic acid. Finally, 800 μL of DMF was added to extract the active components. The samples were vortexed for 1h at room temperature and then stored at -80°C until drug concentrations could be measured using a high-performance liquid chromatography (HPLC) method we described previously (15).

#### 2.6 Cell proliferation assay

Rat aortic smooth muscle cells (RAOSMC; Cell Applications, Inc., San Diego, CA) were grown to 50-60% confluency in 24-well plates. After serum-starvation for 72 hours to achieve cell cycle synchronization, the cells were stimulated with phenol red-free medium containing 10% Fetal Bovine Serum (FBS) and 0.4 mg/mL of either empty pNP (pNP(E)), or quercetin-containing pNP, including pNP(eQ), pNP(eR) or pNP(cQ). Cell proliferation was assessed by following the rate of DNA synthesis, determined as the amount of 5-bromo-2′-deoxy-uridine (BrdU) incorporation (Roche BrDU Labeling and Detection Kit II, Sigma-Aldrich, St. Louis, MO). Briefly, 100 μL BrdU labelling reagent was added to each well and the plates were incubated for 2 h at 37 °C. The medium was aspirated, 300 μL Fixdenat was added, and the plates were incubated for 30 min at room temperature. Next, the Fixdenat was aspirated and 300 μL peroxidase conjugated anti-BrdU antibody was added to all wells, including the background control wells, and the plates were incubated for 90 min at room temperature. The wells were then washed 3 times with 300 μL washing buffer, and 300 μL of substrate were added and allowed to incubate for 2 minutes in dark conditions and at room temperature. Finally, 75 μL 1M H_2_SO_4_ were added to each well, and after rotating for 2 minutes, absorbance was read at 450 nm (reference 690 nm) using a Biotek Synergy microplate reader. Data were expressed as a percent of control cells stimulated with only 10% FBS but with no nanoparticles.

#### 2.7 Adhesion Study Protocol

Because endothelial cells are denuded during angioplasty, smooth muscle cells are the predominate cell type exposed to the balloon to accept pNP containing polyphenols during balloon inflation. Moreover, as explained in the introduction, these cells are typically calcium-laden in PAD. Thus, to model advanced calcified lesion in PAD, RAOSMC were cultured and maintained in a black-walled, clear bottom, tissue-culture treated plates with growth medium compared to calcification medium for two weeks (33). Growth medium contained DMEM with 10% fetal calf serum. Calcification medium contained high glucose (4.5 g/L) DMEM with 10% fetal calf serum, 100 U/mL penicillin, 100 μg/mL streptomycin, 6 mmol/L CaCl_2_, 10 mmol/L sodium pyruvate, 10−6 mol/L insulin, 50 μg/mL ascorbic acid, 10 mmol/L β-glycerophosphate and 10−7 mol/L dexamethasone. Calcification was confirmed using Von Kossa staining (S1 Fig). Next, 10 mg/mL suspensions of pNP(E), pNP(eQ), and pNP(cQ) were diluted in PBS to a final concentration of 2.0 mg/mL. From each suspension 100 µL were placed in wells of the culture plates containing calcified/uncalcified RAOSMC and the cells were incubated at 37°C for 2 hours. We selected 2 hours because pilot observations determined that 2 hours was the minimum amount of time required for pNP to fall to the bottom of the well and adhere. Drug-containing medium was then removed and the cells were subjected to a 100 µL PBS wash before every well was aspirated to dryness. Fluorescence intensity was quantified before and after washing using a Biotek Synergy 2 fluorescence plate reader. Measures of fluorescence detected for cells containing no pNP were used for background correction. In addition, after washing, green fluorescence images of wells were captured on a ZOE Fluorescent Cell Imager (BIO-RAD). Lysis buffer was then placed in wells so that protein levels could be quantified by BCA protein assay. Measures of fluorescence units were normalized to µg protein in each well.

### 2.8. Balloon coating and characterization

#### 2.8.1. Balloon fabrication and ultrasonic coating

A balloon catheter with a 13.8 cm extrusion, a 2.7 FR polycarbonate luer fitting and a 1.25 mm x 10 mm PET over-the-wire balloon was custom manufactured by Interplex Medical, LLC (Milford, OH). Eight balloons were shipped directly to Sono-Tek Corporation (Milton, NY), where they were professionally coated with pNP(eQ) using the following ultrasonic coating method. First, the sample for coating was drawn into a 10 mL syringe, was affixed to a MediCoat BCC coating system and was allowed to reach room temperature. Prior to coating, an atomization test was conducted using a Sono-Tek 48 kHz Accumist nozzle. The material was found to coat flawlessly at low power output. A 3-axis XYZ Gantry System (500 mm x 500 mm x 100 mm), a rotator and the appropriate mounting hardware was interfaced to the system so as to accommodate the balloon catheter. The balloons were inflated and coated using 5, 10, 15 or 20 layers with n=2 balloons coated per group, so that the impact of deposition amount on uniformity and drug loading could be determined. Note that the prohibitive costs of the balloon catheters precluded our ability to test more than 8 balloons.

### 2.8.2. Fluorescence imaging to quantify pNP loading and uniformity of coating

The balloons were affixed on microscopic slides with tape, making every effort to keep them aligned with the center of the slide (deviation of ≤ 3°). Microscopic images were acquired at 4x magnification using a Cytation 3 Image reader (BioTek Instruments Inc, Winooski, VT) in TIFF format, with a 16-bit resolution, both in the visible range and in fluorescent mode. The field of view was panned over in sequential images in order to image the whole balloon in a sequence, and 7-9 images were captured for each balloon. In one case the balloon exceeded the image edges at 4x magnification, so a “top” and a “bottom” image were later combined using the Stitching (34) plugin provided by Fiji (formerly ImageJ) analysis software. The images were acquired with the same imaging parameters (LED intensity=3, integration time=100 ms, camera gain=14), preselected based on the “best” image obtained for a 20 layer-coated balloon, to avoid overly saturating the image brightness of the samples with thinner coatings. As will be apparent in Results, images of one 20 layer-coated balloon (20LYR1) were slightly over-saturated toward the edges of the balloon. However, this did not impact the resulting quantitative measures, as these measurements were performed mainly along the center axis of the balloon. The fluorescent loadings were quantified based on the histograms of two rectangular regions of interest (ROI) per image (Fig 1), each of them 100,000 (500×200) pixels in size (total of 12-14 histograms per coated balloon sample) corresponding to 540,832 µm^2^ (0.54 mm^2^). The ROIs were located along the longitudinal axis as identified using equal distances from the top and bottom edges. The quantification was performed by measuring the mean intensity in each ROI, then averaging the means across all ROIs for a given balloon. Additionally, the overall fluorescence was determined by integrating all brightness values in each histogram and averaging the total brightness across all ROIs for a given balloon. Coating uniformity was determined based on two separate measurements:

**Figure 1.**
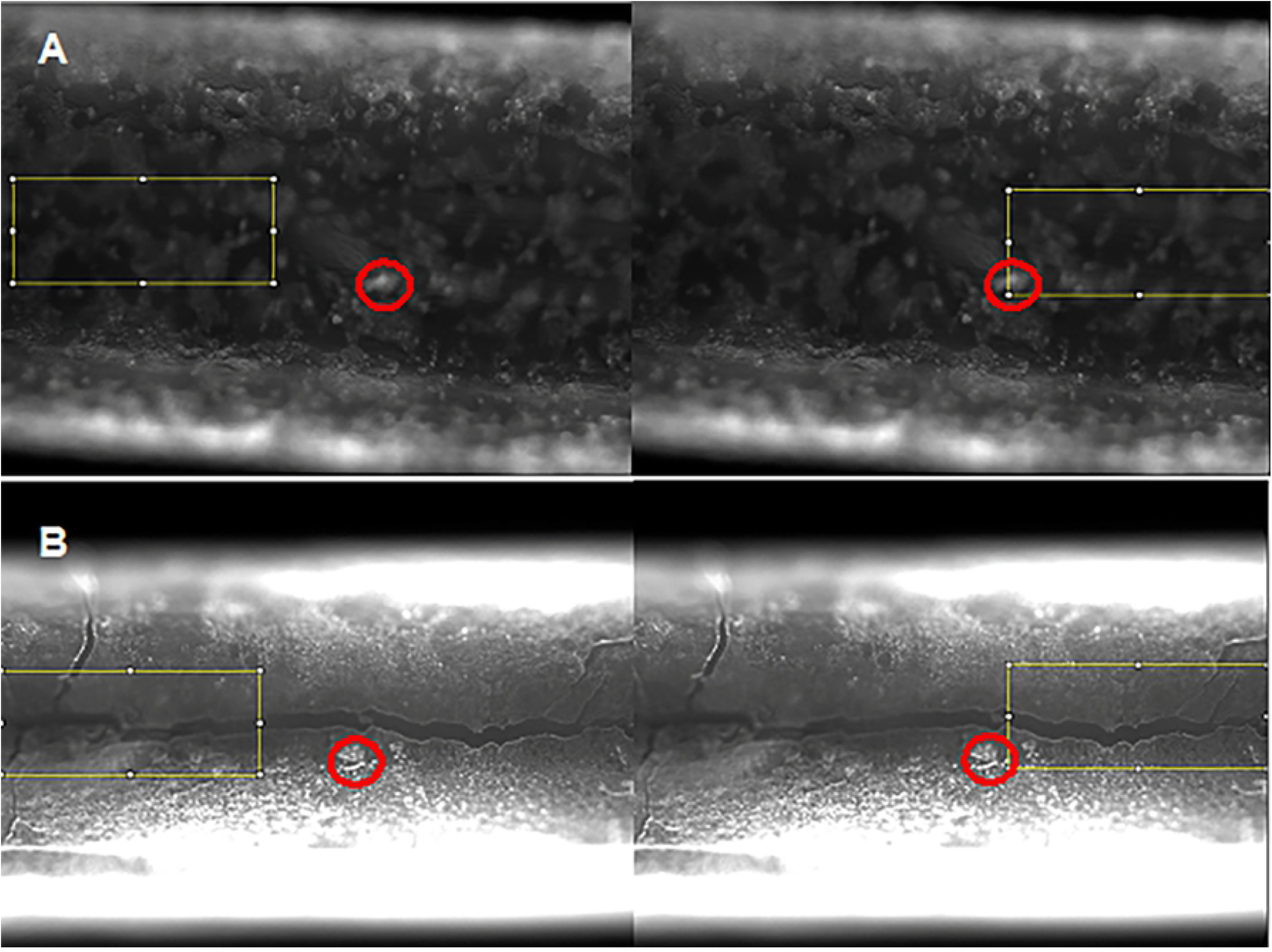
Representative balloon coating images illustrating methods used for assessing uniformity. Red circles indicate points of reference for recording the location of segments examined (yellow boxes). Each picture has a width of 1973 µm and a height of 1457 µm, and the illustration highlights two distinctive areas (left-right) for histogram-based fluorescence analysis. In **A (top row)**, balloon 1 coated with 5 layers (5LYR1), the red circle is 5941 µm from proximal end, and in **B (bottom row)**, balloon 2 coated with 15 layers (15LYR2), the red circle is 5504 µm from the proximal end.

1. The first measurement used the standard deviation of each histogram, with higher standard deviations indicating a less uniform distribution. However, as the images were much “brighter” for the balloons containing higher loading, these values may not be used very reliably to compare balloons possessing differing numbers of layers; i.e. the balloons with fewer layers (thus lower intensities overall) will always have smaller standard deviations compared to the balloons with more layers and larger overall brightness.
2. As an alternative for uniformity of distribution, we also quantified the percent of each histogram area that had brightness intensity within ± 1-SD, which is likely a better indicator of uniformity of distribution, as it indicates how many pixels (or µm^2^) have a brightness of Mean ±SD.

Finally, the fluorescence and brightfield images were stitched together to reconstruct the whole balloon (34), and overlayed for illustration purposes (Fig 2). All image analyses were performed using Fiji software, and corresponding histogram data was exported into Excel for analysis before plotting using GraphPad Prism version 9 Software (La Jolla, CA).

**Figure 2:**
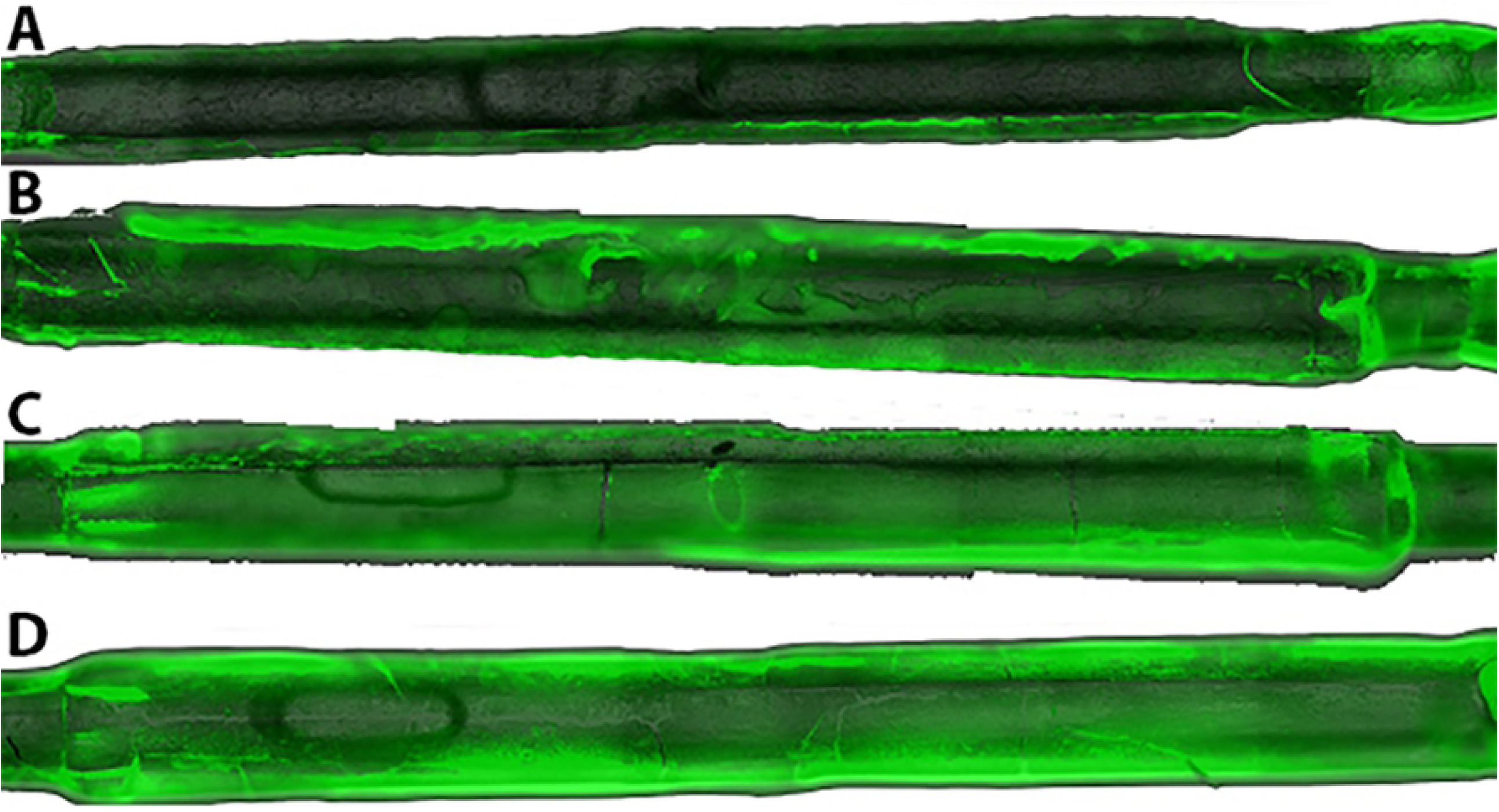
Example of stitched images used to reconstruct the balloons for image analysis. Shown is balloon 2 coated with 5 (**A**), 10 (**B**), 15 (**C**), and 20 layers (**D**) of pNp coating, visible + fluorescent green overlay (at 75% opacity).

#### 2.8.3. Quantification of pNP and drug loading using gravimetric analysis and HPLC

Prior to gravimetric analysis, the balloons were clipped from their catheters and were dried under vacuum for 1 hour. Their weights were measured using a Radwag analytical balance. The coating was then eluted using a 1:1 mixture of 90% acetonitrile: dimethylformamide. The coating suspension was acidified with ascorbic acid, vortexed vigorously and centrifuged. The supernatants were stored at -80°C until HPLC analysis. Finally, the balloons were dried again under vacuum and weighed, so that total coating weights for each balloon could be determined.

## 3. Results and discussion

### 3.1. Nanoparticle characterization

Empty pNP, pNP with entrapped quercetin, pNP with entrapped rhamnazin, and pNP with quercetin covalently attached to PLGA were spherical in shape with a narrow size distribution (Fig 3 and Table 1). The particles ranged in size from 64.9 ± 0.8 nm to 161.9 ± 26.6 nm and were monodispersed (polydispersity index (PDI) < 0.2). The exception was rhamnazin entrapped pNP, which exhibited a PDI of 0.34 ± 0.016 (Table 1). Empty pNP and pNP loaded with polyphenols quercetin and rhamnazin possessed a small positive charge, with zeta potentials of +6.4 - 9.3 mV, while pNPs with covalently attached quercetin possessed a negative charge (zeta potential = -29.9 ± 2.4 mV; Table 1).

**Table 1:** Physical Characteristics of the nanodelivery systems.

|  | Size (nm) | PDI | Zeta potential (mV) |
| --- | --- | --- | --- |
| Empty nanoparticles | 64.9 ± 0.8 | 0.129 ± 0.032 | 6.4 ± 0.3 |
| Entrapped quercetin pNPs | 67.3 ± 1.0 | 0.169 ± 0.003 | 5.9 ± 0.6 |
| Entrapped rhamnazin pNPs | 161.9 ± 26.6 | 0.342 ± 0.016 | 9.3 ± 0.4 |
| Conjugated quercetin pNPs | 106.6 ± 0.8 | 0.050 ± 0.03 | -29.9 ± 2.4 |

**Figure 3:**
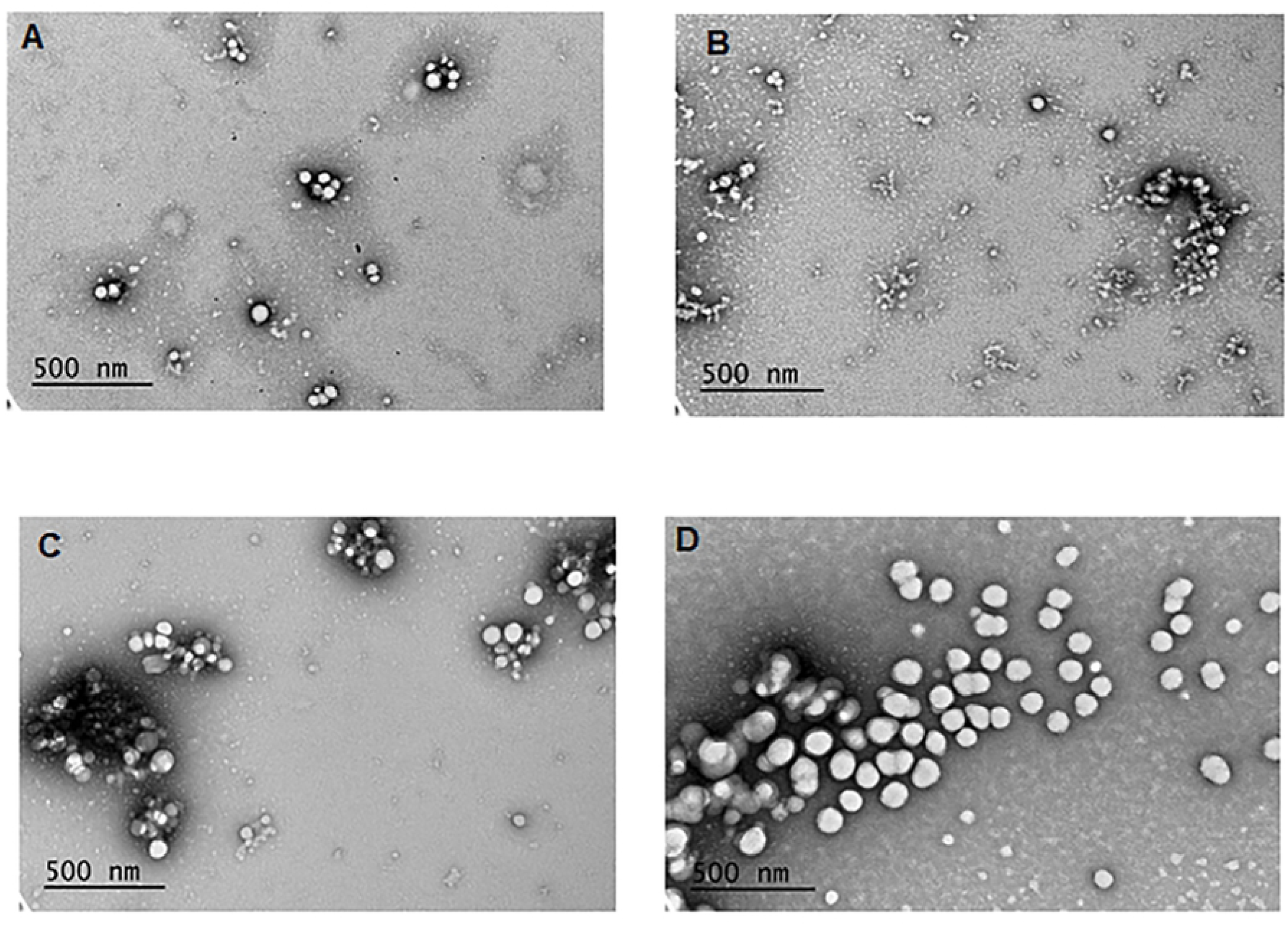
TEM images of (**A**) empty pNP (pNP(E)); (**B**) pNP with entrapped rhamnazin (pNP(eR)) at magnification of 50,000X; (C) pNP with entrapped quercetin (pNP(eQ)); and (D) pNP with covalently attached quercetin (pNP(cQ)) at magnification of 80,000X.

### 3.2. Drug release study

The drug release profile for all 3 entrapped active substances was measured over 6 days. The formulations with entrapped drugs exhibited a burst release within the first day, followed by a more gradual drug release over the remainder of the 6-day period. While entrapped quercetin released rapidly, with 99.7 % of the pNP-entrapped quercetin released by day 3, the release was slightly delayed when more hydrophobic alkylated quercetin (rhamnazin) was used, with 87.7% released by day 3. The covalent attachment of quercetin to PLGA further delayed its release, as indicated by no burst release, only 64.8% release by day 3, and a gradual release over the remaining 3 days of incubation (Fig 4).

**Figure 4:**
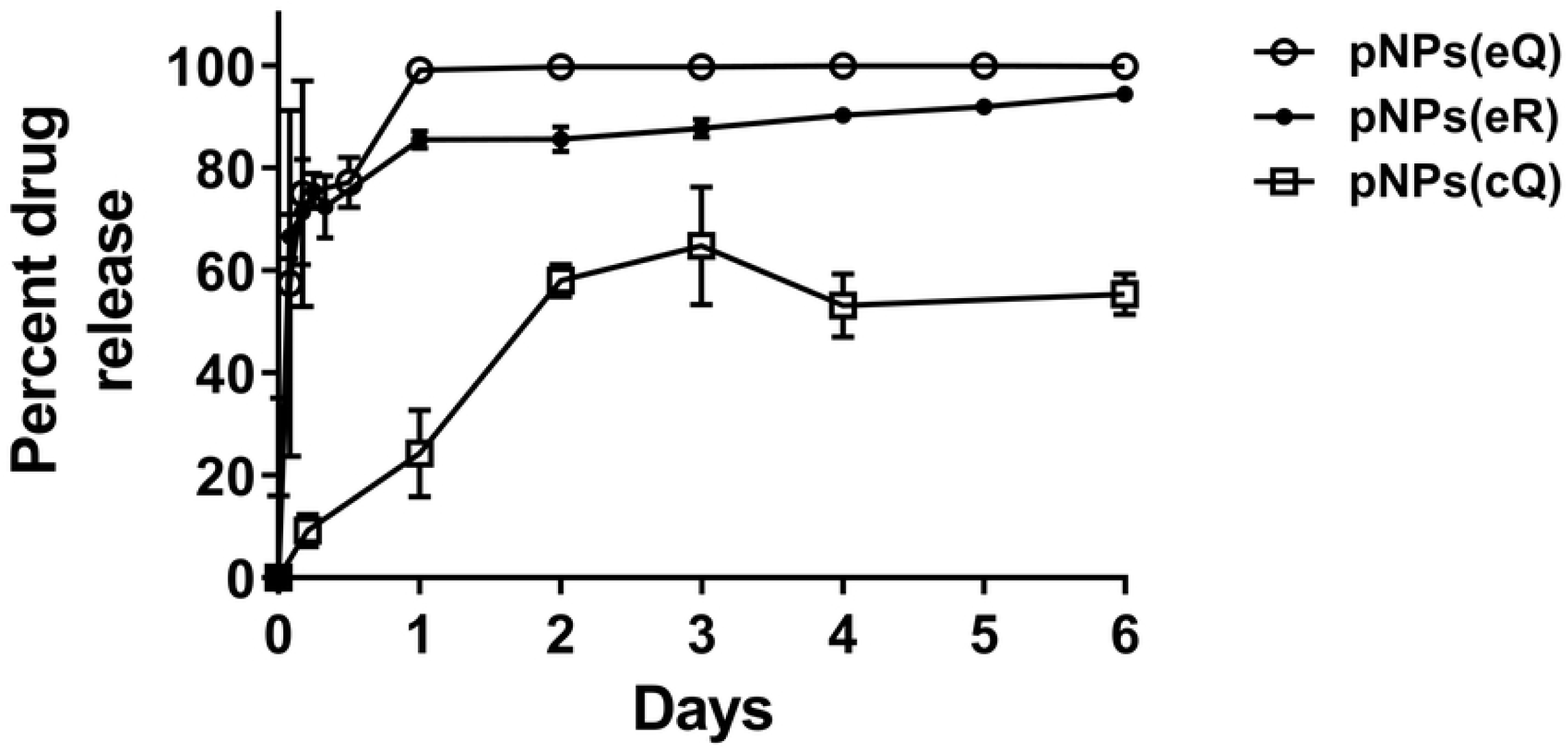
Measures of percent drug release from pNP containing entrapped quercetin (pNP(eQ)), covalently attached quercetin (pNP(cQ)) and entrapped rhamnazin (pNP(cR)). Protracted release was observed mainly for pNPs(cQ). Data are means ± SD for n=3.

### 3.3. Cell proliferation assay

RAOSMC were synchronized, stimulated with 10% FBS ± 0.4 mg/mL empty or drug-loaded pNP for 2 h and rates of cell proliferation were assessed at 24, 48, and 72 hours as relative rates of BrDU incorporation. These relative rates are expressed as a percent of BrDU incorporation assessed for controls cells receiving no treatment. A two-way ANOVA revealed a significant effect of treatment, time and a significant interaction between treatment and time (Fig 5), with all pNP treatments significantly reducing RAOSMC proliferation by 11 to 30% at 24 hours. Note that at this initial time point, even empty pNP – pNP(E) - reduced cell proliferation, though the greatest effect was observed for entrapped quercetin (pNP(eQ)). By 48 hours, however, only the drug-containing particles significantly reduced proliferation and by 72 hours, only pNP covalently modified with Q - pNP(cQ) - maintained its inhibitory effect. Of note, by 72 hours, the empty pNPs exhibited a significant increase in RAOSMC proliferation, although it is unclear whether the 8% increase in proliferation observed for this treatment group and time point is of biologic significance.

**Figure 5:**
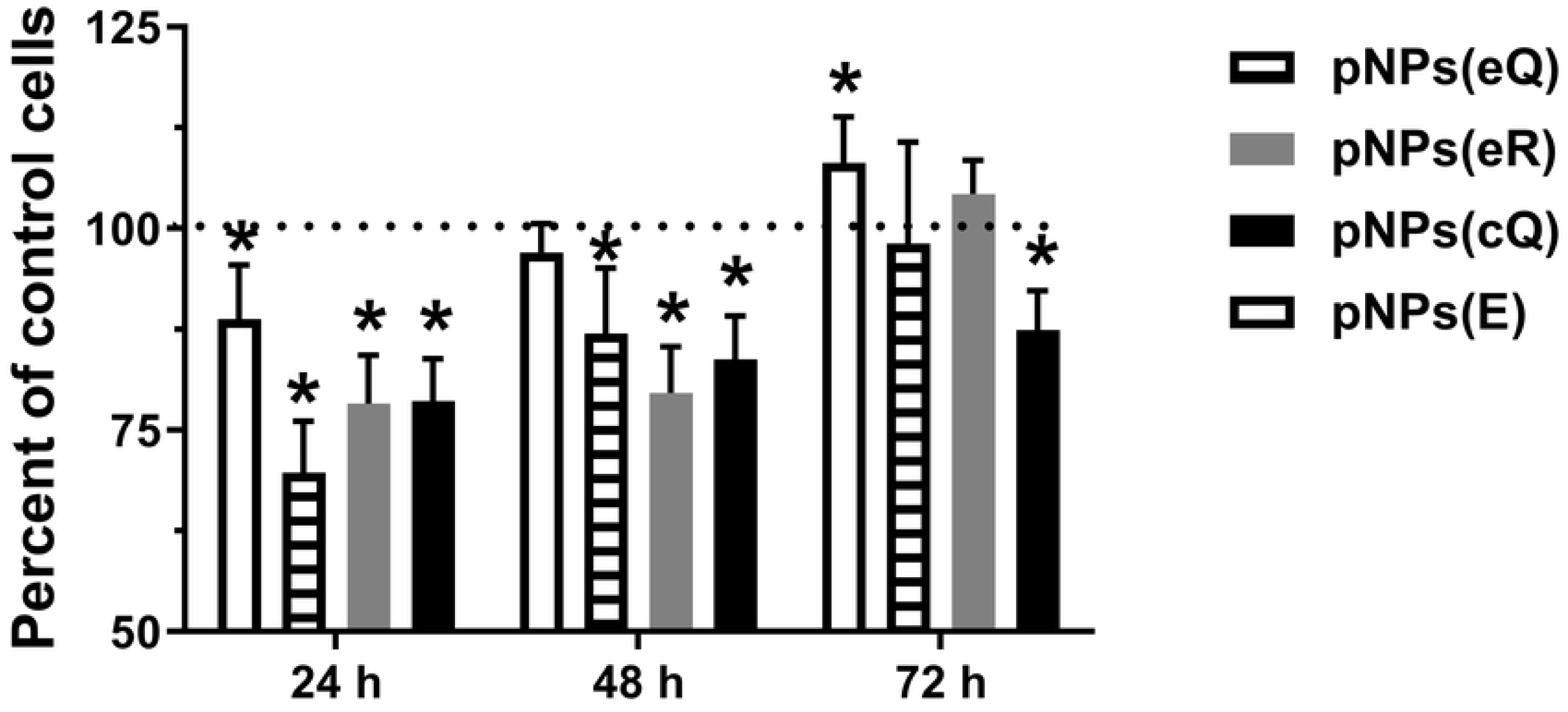
Rat aortic smooth muscle cells loaded for 2 hours with empty pNPs (pNPs(E)), entrapped quercetin (pNPs(eQ)), covalent quercetin (pNPs(cQ)) and entrapped rhamnazin (pNPs(eR)) exhibit reduced rates of cell proliferation at 24, 48 and 72 hours after washing. DNA synthesis was assessed by determining the incorporation of BrDU compared to control cells treated with no pNPs. Data are means ± SD for n=8. Two-way ANOVA revealed a significant effect of treatment. *Indicates significance compared to controls for the same time point, revealed using Dunnett’s post-hoc test. Dotted line represents the response for control cells treated with no pNPs, denoted as 100%.

### 3.4. Measures of pNP adhesion

Zeta potential measures showed that the pNP(cQ) possess a negative, rather than a positive charge. Thus, we hypothesized that upon balloon inflation, these particles would exhibit a reduced ability to bind the negatively charged phospholipid bilayer. However, typically, atherosclerotic arteries in PAD are calcified, with tissues accumulating calcium hydroxyapatite. Calcium hydroxyapatite crystals contain both positive and negative ions and its surface charge is highly dependent upon pH (35) Thus, we further hypothesized that given the ionic nature of calcium hydroxyapatite crystals, the pNP may actually exhibit considerable binding to smooth muscle that has become calcified. To test this hypothesis, we allowed the pNP to adhere to RAOSMCs, with one cohort of these cells cultured under calcification conditions. We used fluorescence imaging to quantify pNP adhesion given the ability of quercetin to fluoresce strongly. Results were that pNP containing Q, including eQ and cQ, exhibited greater fluorescence compared to pNP containing no Q (pNP(E); Figs 6-7). Fluorescence imaging generally supported this finding, except that we noted for cells treated with pNP containing covalently attached quercetin, strong fluorescence was detected in clusters among cells that were calcified (Fig 8). We theorize that perhaps these clusters represent pNP(cQ) binding to calcium hydroxyapatite crystals within the smooth muscle cell cultures.

**Figure 6:**
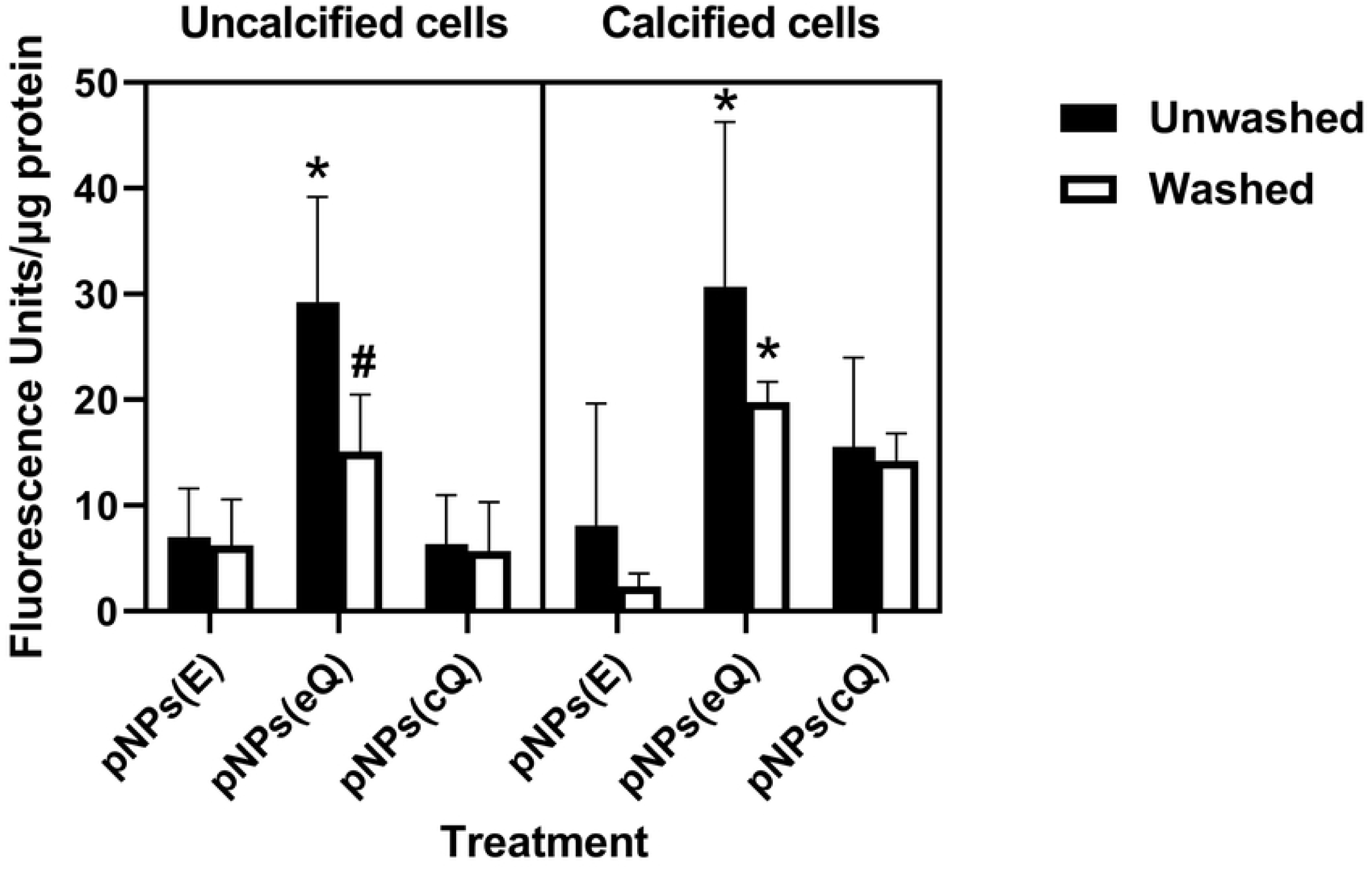
pNPs(cQ) exhibit a reduced ability to bind to rat aortic smooth muscle cells but their binding is resistant to washing and calcification. pNP suspensions at 2 mg/mL were allowed to bind to cells for 2 hours before washing with buffer. Some sets of cells were subjected to a calcification treatment prior to pNP exposure. Green fluorescence determined before and after washing was normalized to protein in the well. Data are means ± SD for n=9. Three-way ANOVA revealed a significant effect of pNP treatment, calcification and washing. *Indicates significance compared to empty nanoparticles (pNPs(E)) for the same cell treatment. #Represents significance compared to unwashed wells for the same pNP treatment.

**Figure 7:**
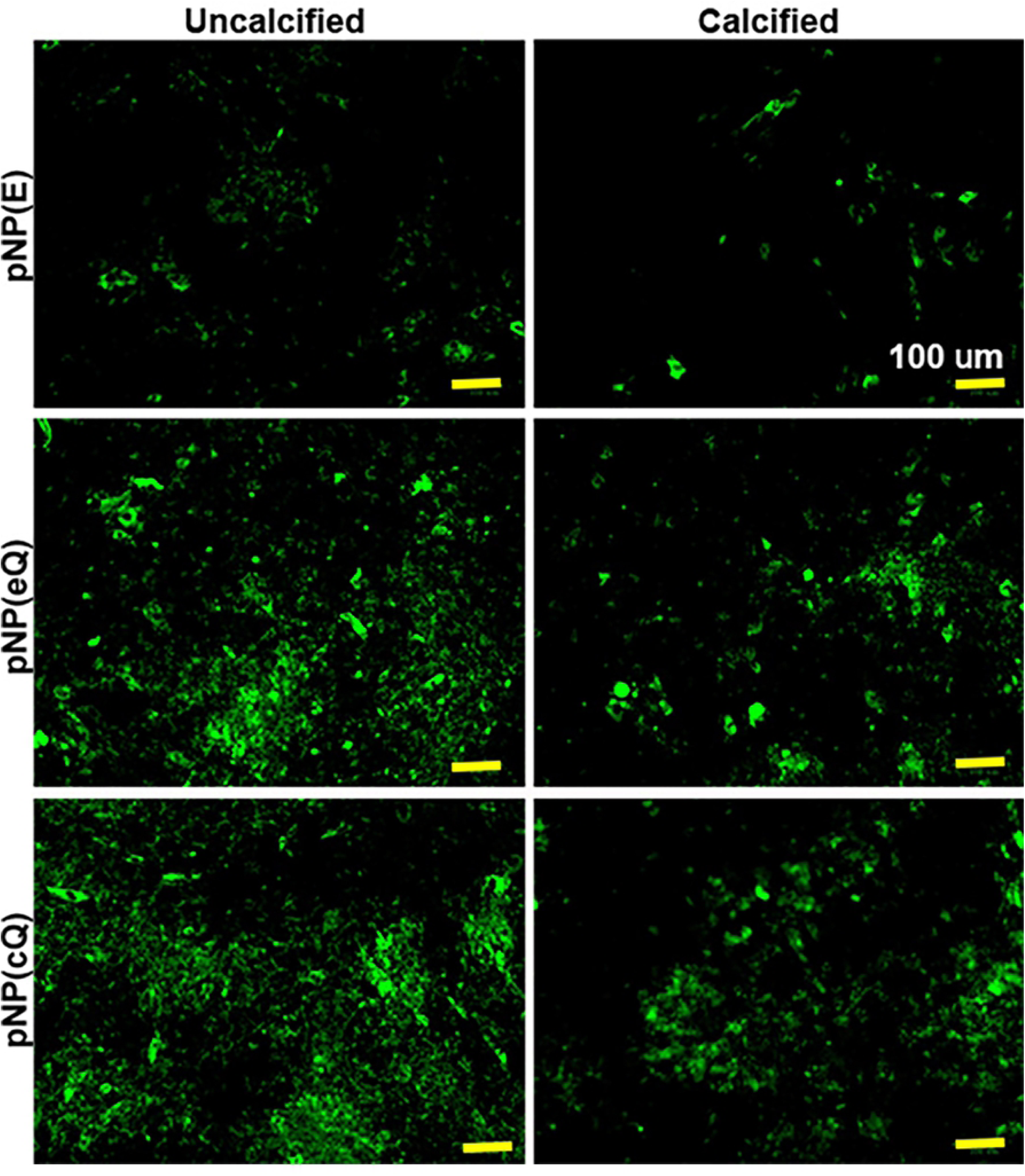
Representative images of green fluorescence within rat aortic smooth muscle cells exposed to pNPs containing quercetin. pNP suspensions at 2 mg/mL were allowed to bind to cells for two hours before washing with buffer. Some sets of cells were subjected to a calcification treatment prior to pNP exposure (right panel). Green fluorescence was imaged after washing. Yellow bar = 100 µm.

**Figure 8:**
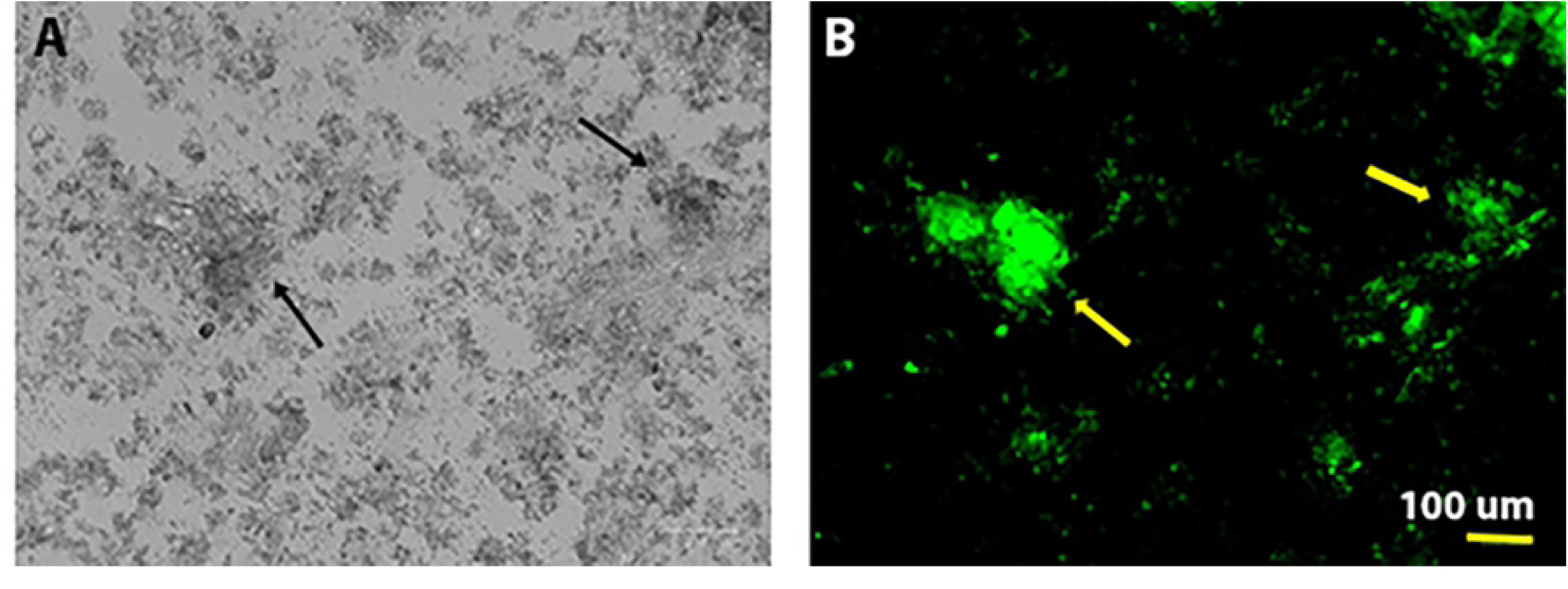
pNPs containing covalently attached quercetin exhibit binding to clusters (indicated by arrows) within calcified smooth muscle cell cultures that were visible in both brightfield (A) and fluorescence (B) images.

### 3.5. Loading based on fluorescence and the uniformity of coating distribution

The results of the image analysis indicate that increasing the number of layers increased the fluorescence intensity of the coating, as expected. Note that balloons were named by denoting 1) the number of layers applied (i.e., 5 layers = 5LYR), followed by 2) balloon sample number (e.g., 5LYR2 = balloon sample 2 coated with 5 layers). There was a clear difference between the samples with 5 layers (5LYR1 and 5LYR2) and the ones with 20 layers (20LYR1 and 20LYR2) (Fig 9A). However, for balloons with an intermediate number of layers (10LYR1-15LYR1), differences in mean brightness were not clearly distinct from one another, even though both of these had a clearly decreased brightness compared to those with 20 layers, and an increased brightness compared to those with 5-layer balloons. These findings are supported by the drug loading data presented in Fig 10, where the balloons with 10 and 15 layers show relatively similar amounts of quercetin loadings. These findings may not necessarily indicate an issue with the coating process, as 10LYR2 and 15LYR2 were found to have good coating uniformity across the balloon surface as indicated by both the standard deviations and percent coverage. Overall fluorescence as determined by integrating brightness values over the whole ROIs yielded similar results as the mean values and thus, are not presented here.

**Figure 9.**
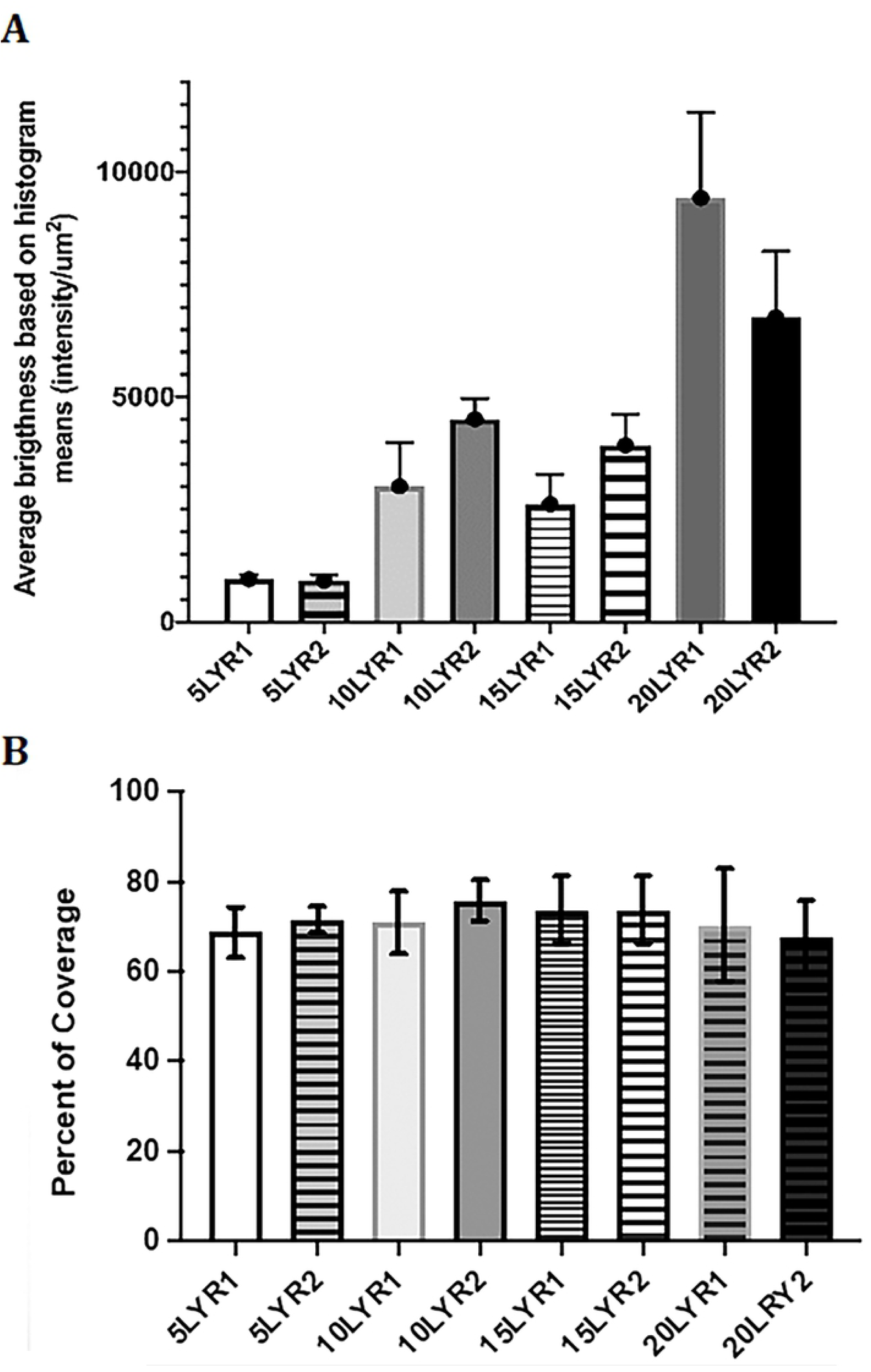
Fluorescence imaging revealed that ultrasonic coating with pNPs entrapping polyphenols yields uniform coatings. A) Overall mean fluorescence and corresponding standard deviations for n=8 samples (balloons ultrasonically coated). Data illustrated in the graph represent mean fluorescence ± SD (in unit of brightness per µm^2^). Maximum brightness for a 16-bit image is 65535, corresponding to 12117.42 per µm^2^. B) Percent of balloon area that has pixels with fluorescence intensity within a ± 1-SD of the mean fluorescence. Data represent means ± SD for n=8 samples. Higher value indicates better uniformity.

**Figure 10.**
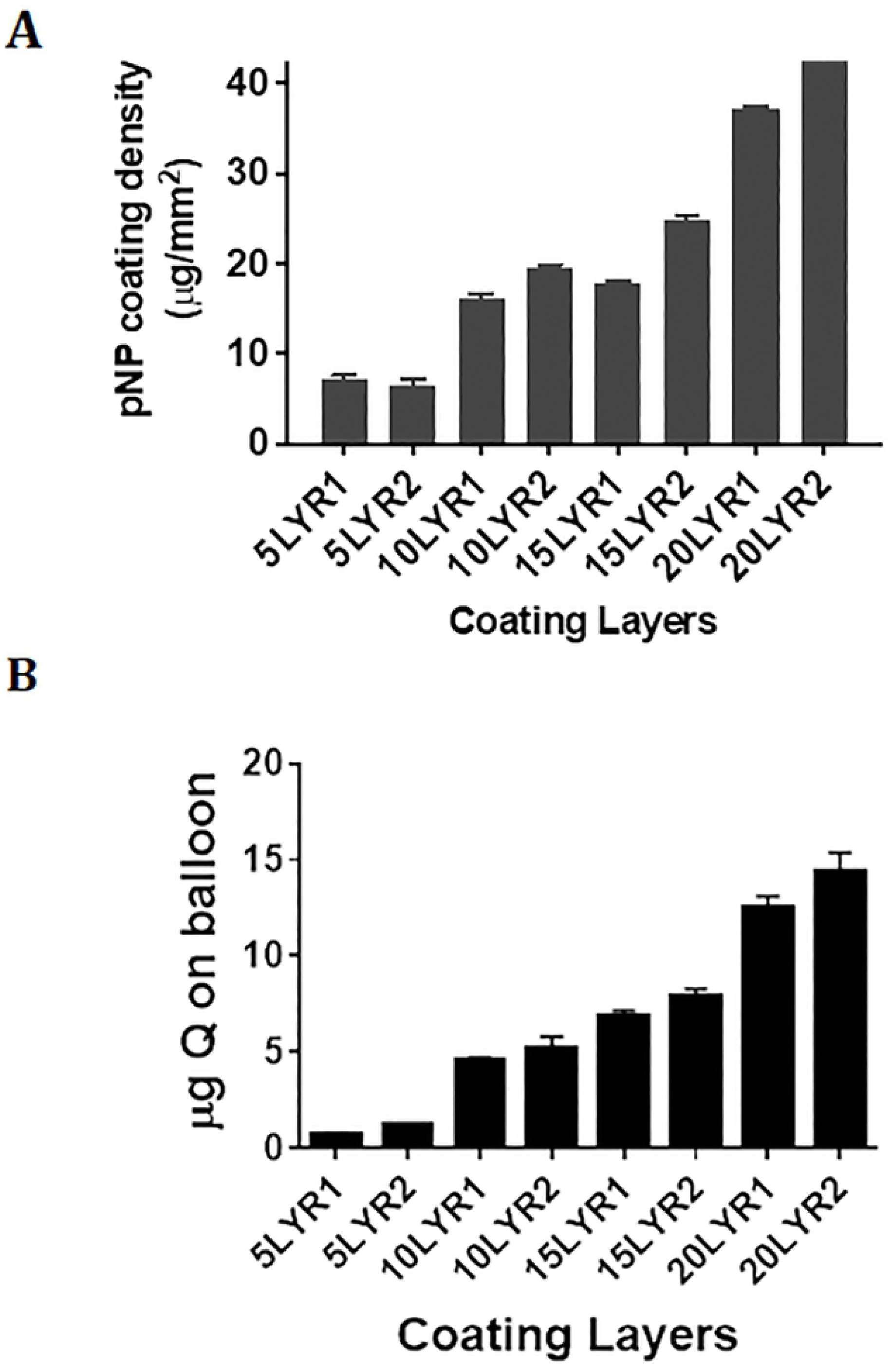
Amount of quercetin (Q) and quercetin nanoparticles (pNPs) in balloon coating. A) pNPs were eluted from the balloons using organic solvent and pNP load was determined gravimetrically. Total loading in µg was normalized to balloon areas. Data represent means ± SD for 5 measures per balloon. B) pNPs were eluted from the balloons using organic solvent and the Q content was determined using HPLC. Data represent means ± SD for n=3-4 replicate measures/balloon.

Standard deviations for the histograms (Fig 9A) suggest that the uniformity of coating deposition decreases with an increasing number of layers deposited. However, these findings may be biased by the fact that balloons with fewer coating layers would have much lower overall brightness and thus, smaller standard deviations associated with those mean values. To compensate for differences in standard deviations due to differences in the magnitude of overall brightness, standard deviations were normalized to the mean of each histogram (Table 2). This additional analysis indicates that two of the balloons (10LYR1 and 15LYR1) exhibited a deviation of more than 25% of the mean value, which may be indicative of a lower uniformity of coating compared to the other samples.

**Table 2:** Average fluorescence brightness, absolute standard deviations, and normalized standard deviations for the coated balloons (max possible brightness = 12117.4/µm2).

| | Average brightness<br>(intensity/$\mu\text{m}^2$) | Absolute STD<br>(intensity/$\mu\text{m}^2$) | Normalized STD<br>(% of mean) |
| --- | --- | --- | --- |
| 5LYR1 | 955.38 | 108.16 | 11.3% |
| 5LYR2 | 908.64 | 137.25 | 15.1% |
| 10LYR1 | 3029.37 | 975.13 | 32.2% |
| 10LYR2 | 4502.90 | 458.34 | 10.2% |
| 15LYR1 | 2547.79 | 661.59 | 26.0% |
| 15LYR2 | 3919.94 | 694.69 | 17.7% |
| 20LYR1 | 9422.61 | 1905.54 | 20.2% |
| 20LYR2 | 6787.45 | 1467.44 | 21.6% |

The second histogram-based measurement of coating uniformity was determined by quantifying the percent area in each histogram that has pixels with brightness (*i*.*e*. fluorescence) within ± 1-SD of the mean value. This measurement should be independent of absolute pixel brightness in a given ROI. Thus, this value can be used more reliably to compare uniformity between samples with differing numbers of layers (Fig 9B). Based on these measurements, the percent area covered ranged from 67.6 to 75.8%, which can be considered from good to excellent coverage or uniformity, based on a uniformity scale of: <55% poor, 55-60% moderate, 60-70% good, 70-75% very good, 75-80% excellent, >80% outstanding, with outstanding very rarely occurring in normal image processing of spray-type coatings (a 100% value would indicate all pixels in the ROI having the exact same value, which would be nearly impossible to achieve).

Several of the balloon samples showed cracking of the fluorescent layers, clearly visible in the fluorescent images which may have an unquantified influence on the results of the image analysis, but based on the visual inspection of the images, these were relatively sparse and otherwise small overall. As the coating process occurred in a different location than the fixation on the slide and subsequent imaging, it cannot be ascertained if the cracks are a result of the coating process itself or an artifact introduced by the maybe too rapid drying after coating, or by handling during transport and slide fixation. As the samples with the smallest number of layers (5LYR1 and 5LYR2) did not exhibit any visible cracking, it seems that this phenomenon occurs only for thick layers, which, upon drying, are more prone to cracking.

### 3.6. Loading of pNP and quercetin, assessed using gravimetric analysis coupled to HPLC

Gravimetric analysis mirrored the results of the fluorescence analyses. pNP coating weights increased nearly linearly with increasing numbers of layers, although coatings with 10 and 15 layers contained more similar amounts of deposited pNP compared to other groups (Fig 10A). In total, 0.26-1.5 mg of pNP were successfully applied through 5-20 coating layers, respectively (not shown). Adjusting for the surface area of the balloon, this amounted to 7-40 µg/mm^2^ (Fig 10A).

HPLC analysis of pNP eluted from the balloons revealed a more linear increase in quercetin levels as the coating layers were increased, with total quercetin loading ranging from 0.8-14 µg through 5-20 layers, respectively (Fig 10B).

## 5. Conclusions

Peripheral artery disease (PAD) is an inflammatory disease primarily caused by atherosclerosis, which gradually narrows the arterial lumen. Revascularization is considered the first line therapy for symptomatic obstructive PAD (10, 36). Catheter-based percutaneous interventions are an enduring relief for arterial obstruction (36) and are considered the primary method for revascularization (36-38). Restenosis is defined by a reduction in the diameter of the vessel lumen after angioplasty (39). Much research and commercialization effort has been devoted to manufacturing device technologies targeting restenosis (40). The use of polymeric or metallic stents provides better acute results, but these improvements arise at the expense of increased vessel injury (36, 41, 42), with stents commonly resulting in increased risks of thrombosis and stent fracture (43, 44). The need to address the associated risk that comes with stenting led to non-stent-based local drug delivery. Drug-coated balloons are alternative approaches in which the balloon is coated with a thin, active substance surface layer (Byrne et al., 2013). Delayed healing along with vascular toxicity of the anti-proliferative agents applied to the balloon’s surface was observed in animal studies after DCB angioplasty (36). In our own prior studies, a nanoparticle delivery system was designed to provide an alternative treatment for PAD, using polyphenols with high therapeutic indices as alternatives to the anti-proliferative agents in commercial products (15). Similar coatings releasing quercetin and resveratrol from drug eluting stents demonstrated outstanding effects in reducing VSMC proliferation, platelet activation and inflammation, while promoting re-endothelization (45, 46). The cationic characteristics of the pNP were provided by addition of a cationic Eudragit RL100 polymer during pNP synthesis. By adjusting the amount of positive charge on the system, the pNP were designed to be biocompatible and biodegradable and proved to meet the specification ideal for cellular uptake and maintaining a continued period of release. The PLGA nanoparticles with pNP(eQ), pNP(eR), as well as quercetin conjugated to PLGA (pNP(cQ), were developed at a size range of 101 nm. All polyphenols were entrapped separately in PLGA pNPs to allow for their comparison. Similar to prior experiments, entrapped quercetin released rapidly in the first 24 hours except that this time the active substance was entrapped separately in pNPs not together with RESV in its methoxylated form (15). However, covalent attachment of quercetin delayed its release as indicated by no burst release and a more protracted profile. The methoxylated derivative of quercetin (rhamnazin) with increased hydrophobicity provided a slightly more sustained release of quercetin, although was not as protracted as pNPs possessing covalently-attached Q. In the latter case, release was sustained for a total of 6 days, which is beneficial since vascular healing, as well as the cellular events contributing to restenosis, begin within the first 7 days (47). In this experiment an ultrasonic coating method was used that allowed our pNP entrapment system to generate a uniform coating. This coating technique will hopefully minimize non-specific release of drug into the blood and enhance the long-term retention of drug within vascular tissue, but such specifications will be addressed in future animal experiments.

In summary, a key parameter for a successful DCB is delivery of therapeutic levels of drug at biologically appropriate time points within a critical time window after endovascular intervention. The synthesized PLGA-based pNP system proved to be biocompatible with a size range required for endocytosis and provided an extended period of release. Importantly, brief application with pNPs containing covalently-attached Q demonstrated an ability to reduce VSMC proliferation at least through 72 hours. Studies utilizing a balloon angioplasty model in small animals aimed at testing the pharmacokinetics of drug delivery to the vascular wall will be required for further development.

## Acknowledgments

This work was funded by the National Heart, Lung & Blood Institute, National Institutes of Health, R41 HL142403 and the Louisiana Board of Regents Industrial Ties Research Subprogram. The authors would like to acknowledge partial support from the LSU Biological and Agricultural Engineering Department at LSU AgCenter and USDA NIFA Hatch program (LAB #94443 and LAB #94513). Published with the approval of the Director of the Louisiana Agricultural Experiment Station as manuscript #2022-232-36870.

## Disclosures

CAE, CMS and TRD have intellectual property related to the work presented in the manuscript. TRD is a co-founder of a biomedical company aimed at developing angioplasty balloon coatings.

